# A reaction norm for flowering time plasticity reveals physiological footprints of maize adaptation

**DOI:** 10.1101/2024.07.10.602692

**Authors:** Justine Drouault, Carine Palaffre, Emilie J. Millet, Jonas Rodriguez, Pierre Martre, Kristian Johnson, Boris Parent, Claude Welcker, Randall J. Wisser

## Abstract

Understanding how plant phenotypes are shaped by their environments is crucial for addressing questions about crop adaptation to new environments. This study focused on analyzing the genetic variability underlying genotype-by-environment interactions and adaptation for flowering time in maize. We present a physiological reaction norm for flowering time plasticity (PRN-FTP), modeled from multi-environment trial networks and decomposed into its physiological components. We show how genotype-specific differences in developmental responses to temperature fluctuations condition differences in photoperiod perceived among genotypes. This occurs not only across but also within common environments, as the perception of photoperiod is altered by variation in rates of development and durations for becoming sensitized to photoperiod. Using a new metric for envirotyping sensed photoperiods for maize, it was found that, at high latitudes, different genotypes in the same environment can experience up to hours-long differences in photoperiod. This emphasizes the importance of considering genotype-specific differences in the experienced environment when investigating plasticity. Modeling the PRN-FTP for globally representative breeding material showed that tropical and temperate germplasm occupy distinct territories of the trait space for PRN-FTP parameters. Placed in the historical context of maize, our findings suggest that the geographical spread and breeding of maize was mediated by a specific modality of ecophysiological adaptation of flowering time. Our study has implications for understanding crop adaptation and for future crop improvement efforts.

**Article Summary:** Maize adjusts its flowering time across environments using genetic pathways tuned to cues like temperature and day length. This study models how different varieties perceive and respond to these factors, revealing that genotypes can perceive the same environment in distinct ways. By disentangling the ecophysiological basis of genotype-environment interactions, the research explores maize diversity and highlights how adaptation and breeding have shaped distinct strategies for flowering time regulation in tropical and temperate varieties.

## Introduction

Plants exhibit vast phenotypic diversity, the principal axis for adaptation and evolution. The classic formalism *P* = *G* + *E* + *G* × *E* describes how phenotypes are determined by combinations of genetic (*G*) and environmental (*E*) effects. Environmental effects result from varying external conditions that lead to distinct phenotypic outcomes for the *same* genotype, a phenomena known as plasticity (Bradshaw, 1965). As 21^st^-century climate markedly shifts the environmental dimension to new frontiers, it is an important and opportune time to investigate plasticity (*E*) and genetic variation in plastic responses (*G* × *E*). For crop improvement, harnessing plastic responses to shifts in climate conditions can facilitate the adaptation of crops into new territories or to alternative cropping systems (Rising and Devineni, 2020; Sloat *et al*., 2020; Zabel *et al*., 2021). Thus, approaches for modeling *G* × *E* that are rooted in the understanding of plasticity represent a promising direction for agricultural research (e.g., Millet *et al*., 2019; Cooper *et al*., 2020; van Voorn *et al*., 2023).

The functional relationship that defines a pattern of plasticity across an environmental gradient is known as a reaction norm, with parameters that quantify the direction and magnitude of phenotypic change (Roof, 1997). The reaction norm provides a foundation for modeling phenotypic responses to the environment. However, separate disciplines have conceived of reaction norms in different ways. A common approach in quantitative genetics uses the *Finlay–Wilkinson* statistical model applied to multi-environment trials (METs) (Finlay and Wilkinson, 1963), in which the phenotypic values of individual genotypes are regressed on an index defined by the mean values of all genotypes per environment. In this framework, phenotypes are typically for final characteristics measured at the whole-plant scale that are targets for selection, such as flowering time, plant height, or yield. This method has provided a convenient generalization for modeling reaction norms from METs where large collections of genotypes can be simultaneously tested across different environments, enabling a widescale analysis of *G* × *E* (Jarquín *et al*., 2014; Kusmec *et al*., 2017; Guo *et al*., 2023). Clearly, however, the environment is abstractly defined and confounded with latent environmental effects where mechanisms underlying *G* × *E* become elusive, resulting in a lack of biological interpretability. In contrast, focused on phenotyping of finer-scale traits in controlled settings, ecophysiologists have characterized norms of reaction for physiological responses to specific environment variables, such as the non-linear response of growth and development to temperature (Parent *et al*., 2010; Parent and Tardieu, 2012). Embedding formalisms for such reaction norms in crop growth models allows an upscaling to the phenotypes for final characteristics. This is modeled as a function of genotype-specific parameters and environmental conditions, with outcomes emerging from physiological mechanisms that dynamically regulate plant development in response to fluctuating conditions across time (Muller and Martre, 2019). This approach provides biological perspective, but studying broad diversity of *G* × *E* is challenged by experimental constraints to measuring the parameters of physiological reaction norms. In our study, we hypothesized that parameterizing a physiological reaction norm from MET data would allow us to analyze the diversity of flowering time plasticity in maize (*Zea mays* subsp. *mays*) and provide unique insights into maize adaptation.

Physiological responses to multiple environmental factors modulate development which affects flowering time, particularly temperature and photoperiod (Simpson and Dean, 2002; Colasanti and Coneva, 2009; Blümel *et al*., 2015; Lee *et al*., 2021). The influence of temperature results in a speeding or slowing of development, while, for photoperiod sensitive genotypes, the influence of photoperiod results in the suppression of reproductive transition and a delay in flowering time (this occurs under long daylengths for short-day species and short daylengths for long-day species). The response to temperature occurs throughout a plant”s life cycle, while the response to photoperiod is growth-stage specific. This is a crucial distinction for the work presented here. In maize, a facultative short-day (SD) species, the stage at which plants become sensitized to photoperiod is not directly observable, but based on photoperiod-swap experiments has been found to occur immediately before tassel initiation (reproductive transition) if plants are grown specifically under SD conditions (Kiniry *et al*., 1983*b*; Bassiri *et al*., 1992). In SD conditions, tassel initiation is not repressed by photoperiod and temperature is the primary driver of development time and determinant for when a genotype reaches the sensitized stage. Above a critical photoperiod, under long-day (LD) conditions, tassel initiation in photoperiod sensitive genotypes is delayed beyond the timing at which they reach their sensitized stage, with a time delay that depends on genotypic variation in the degree of photoperiod sensitivity in addition to temperature fluctuations during the period of delay. Thus, as physiological responses to temperature fluctuations across the development cycle interact with genotypic variation in rates of plant development and durations of the reproductive transition, different genotypes can reach the sensitized stage on different days, and therefore perceive different photoperiods, even within the same environment. For photoperiod sensitive genotypes, the magnitude of response to photoperiod is an additional source of plasticity that varies genetically, culminating in the formation of *G* × *E* interactions in flowering time across environments. We introduce these first-principles expectations for *G* × *E* interactions in flowering time, but we are not aware of studies that have considered this in the construction of reaction norms from MET data.

The timing of reproductive transition (and consequently flowering) is controlled by a genetic regulatory network (GRN) that integrates both the exogenous and endogenous signals (Song *et al*., 2015). A reference topology of this GRN in maize consists of multiple pathways including the circadian clock, photoperiod, autonomous, aging, and gibberellic acid pathways (Dong *et al*., 2012). However, the complete topology and context-dependent expression of the GRN is far from being fully understood (Minow *et al*., 2018; Stephenson *et al*., 2019). A key distinction exists in the way the GRN functions between LD and SD environments, with daylength-dependent and daylength-independent responses for photoperiod sensitive and insensitive genotypes. For example, the circadian clock pathway plays a central role in the transduction of signals to the other pathways, with a direct link between external light conditions and the photoperiod pathway (in both LD plants like Arabidopsis – Sawa *et al*., 2007; Sawa and Kay, 2011; and SD plants like maize - Bendix *et al*., 2013). Under LD conditions, transduction of light perception by the circadian clock to the photoperiod pathway represses production of the *FT*-derived florigen signal for reproductive transition in photoperiod sensitive genotypes of maize—flowering time is delayed (Meng *et al*., 2011). In contrast, the mechanism controlling floral transition is configured differently in insensitive genotypes of maize, with recent research implicating separate clock components that interact with the autonomous pathway (Minow *et al*., 2018). Less is known about the determination of critical thresholds for photoperiod sensitivity. A gating mechanism involving the circadian clock and photoperiod pathways has been described in rice (a facultative SD species like maize; Itoh *et al*., 2010; Nemoto *et al*., 2016), with additional findings suggesting a conditional link involving the autonomous pathway described in maize (Matsubara *et al*., 2008; Park *et al*., 2008).While molecular genetic studies continue to elucidate the intricate dynamics of cross-talk and signal transduction by the GRN, a challenge is to relate this to functional diversity at the whole plant level. Thus, new approaches to characterizing the wider scale variation in ecophysiological plasticity for flowering time are needed.

This study focused on the modeling and analysis of a physiological reaction norm for flowering time plasticity (PRN-FTP) that integrates genotype-specific sensing and responses in growth and development to fluctuating temperatures and photoperiods, ultimately resulting in *G* × *E* for flowering time. Building on previous work (Kiniry *et al*., 1983*a*,*b*; Choquette *et al*., 2023), the PRN-FTP was formulated as a bilinear, threshold-dependent function that relates thermal time from crop emergence to flowering with a novel envirotyping index for the daylength sensed (*DL*^s^) by individual genotypes in each environment. We envirotyped *DL*^s^ across field environments in the Northern Hemisphere to understand how photoperiod perception varies among genotypes both within and across environments, and whether genotype-specific values for *DL*^s^ should be used for modeling the PRN-FTP. We further demonstrate how analyzing the PRN-FTP as a bilinear function allows the decoupling of different components of flowering time variation: (i) flowering time *per se*; (ii) critical photoperiod; and (iii) photoperiod sensitivity; with genotype-specific parameter values estimated from MET data.

Finally, PRN-FTP parameters may be used to understand how flowering time plasticity varies among population groups or mediates adaptation to novel environments, and how this in turn alters flowering time plasticity for the future. Historical varieties of maize that spread from their tropical origins into temperate zones required adaptations to cooler climates with longer photoperiods, with studies implicating the central role of flowering time (Tenaillon and Charcosset, 2011). For instance, several flowering time genes in the GRN for flowering time have been linked to the post-domestication spread of maize (e.g., *Vgt1* – Ducrocq *et al*., 2008; *ZmCCT10* – Hung *et al*., 2012; Yang *et al*., 2013; *ZCN8* - Guo *et al*., 2018; *ZmELF3*.*1* – Zhao *et al*., 2023). Here, we show that the trait space relating variation in the PRN-FTP parameters flowering time *per se* and photoperiod sensitivity putatively reflects how adaptation and selection history has shaped differences in flowering time plasticity between breeding pools of modern maize.

## Materials and Methods

A general guide to the study design and use of datasets 2-4 is presented in Fig. 1.

**Figure 1.**
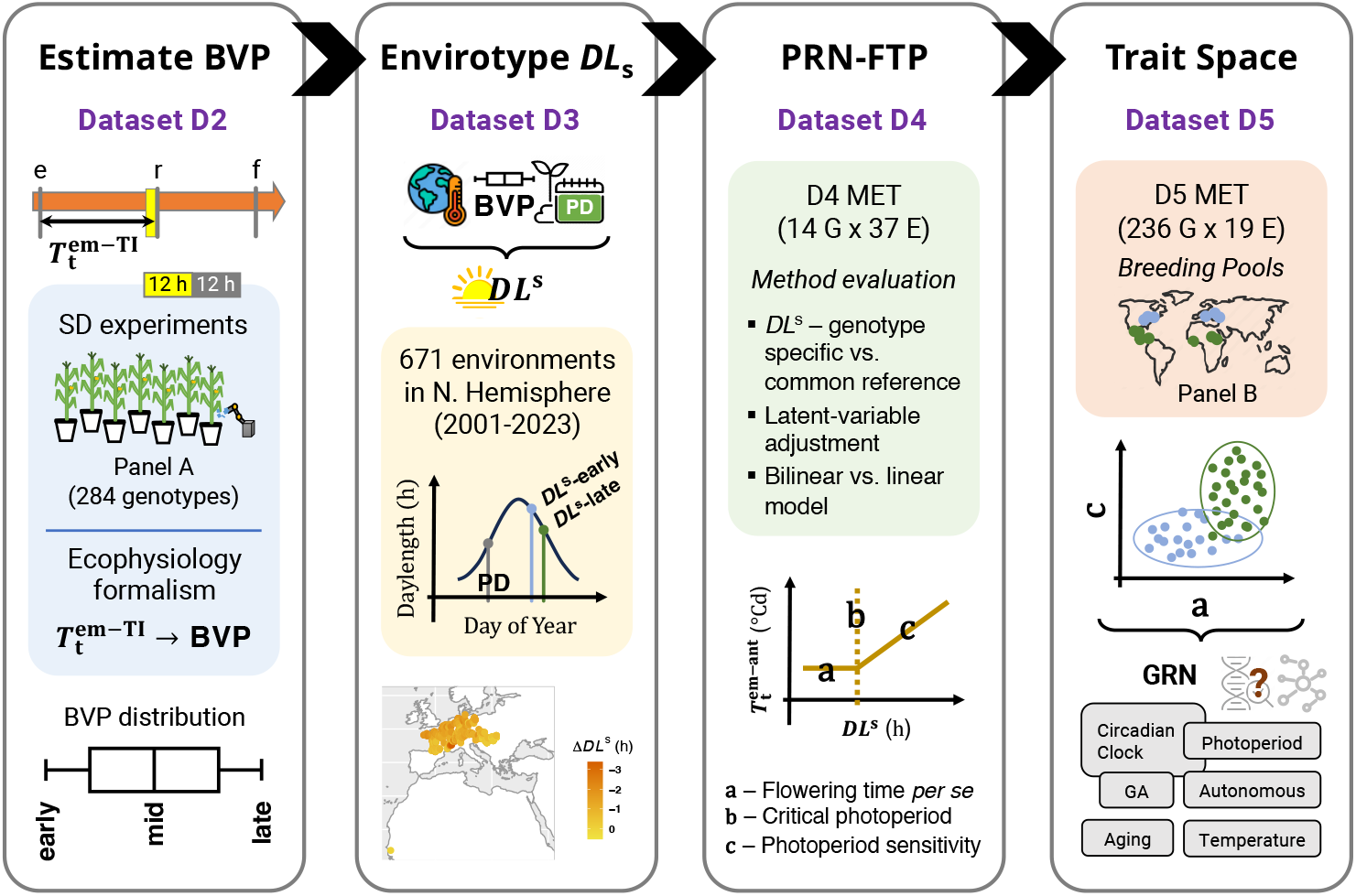
Graphical guide to the study. Each panel portrays the main parts of the study performed with Datasets 2-4 (Dataset 1 was used to evaluate the choice of a temperature response function for calculating thermal time, not shown): (i) The estimation of basic vegetative phase (BVP) follows from prior research showing that the photoperiod-sensitized stage is reached just prior to the thermal-time required for reproductive transition 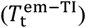 plants grown under SD conditions. This is depicted by the filled arow marked with the timing of emergence (“e”), reproductive transition (“r”), and flowering (“f”). The orange shading of the arrow represents that plants sense temperature throughout their life cycle, while the yellow box represents that photoperiod begins to be sensed at a particular time. Data from short-day (SD) experiments were used to estimate the BVP distribution for 284 genotypes in panel A. (ii) The temperature data from maize fields, BVP reference values for early, mid, and late maturing genotypes, and planting dates (“PD”) were used to envirotype the daylength sensed (*DL*^s^) by each maturity class in 671 environments. The figure in the shaded region depicts how *DL*^s^ can vary between early and late maturing genotypes that become sensitized to photperiod on different days of the year (this can vary due to fluctuations in temperature and rates if photoperiodic change). This was used to characterize inter- and intra-environmental variation in *DL*^s^ for maize in the Northern Hemisphere. (iii) The physiological reaction norm for flowering time plasticity (PRN-FTP) was modeled using MET data on 14 genotypes in 37 environments. An evaluation of the methodologies for modeling the PRN-FTP was performed as noted in the shaded area. The figure portrays the PRN-FTP modeled as a bilinear function with coefficient estimates for flowering time *per se* (a, units °C), critical photoperiod (b, units: h), and photoperiod sensitivity (c, units: °Ch^-1^). (iv) The PRN-FTP was modeled as a linear function (LD-SD method, see text) using MET data on 214 genotypes (panel B) in 19 environments. The trait space defined by flowering time *per se* (a) and photoperiod sensitivity (c) was used to infer about historical adaptation in maize. The map in the shaded area portrays genotypes from temperate (blue) and tropical (green) breeding pools of maize comprising panel B. The trait-space plot reflects a finding of this study, showing that tropical and temperate breeds occupy different territories of the space. Future research on the genetic dissection of the separate parameters may be used to relate ecophysiological scale variation to pathways in the gene regulator network (GRN) for reproductive transition and flowering time.

### Datasets

Summarized in Table S1, datasets (“D”) from multiple projects and experiments on maize were assembled to: (D1) select a temperature response function for thermal time estimation; (D2) develop a predictor function for *DL*^s^; (D3) envirotype *DL*^s^ for diverse genotypes in fields across the Northern Hemisphere; (D4) demonstrate methods for modeling a physiological reaction norm for flowering time plasticity (PRN-FTP); and (D5) compare the trait space of parameters estimated from the PRN-FTP in a diversity panel constituting separate breeding pools of maize. A brief description of the datasets used for these five objectives are presented first, followed by methods and other details in subsequent sections. The current study focused on days to anthesis (male flowering) as the final characteristic.

(D1) This dataset contains 24 hybrid genotypes measured for days to anthesis in 25 to 26 field experiments, spanning latitudes from 30.5371 °N to 44.2089 °N (File S1). For each environment, the sowing date, field coordinates, and best linear unbiased estimates (BLUEs) for calendar days from sowing to anthesis were obtained from McFarland *et al*., 2020 and Rogers *et al*., 2021.

(D2) This dataset contains a collection of 284 inbred genotypes (panel A) with trait data combined from two experiments. Emergence time and phyllochron were measured under well-watered conditions in a greenhouse phenotyping platform at the LEPSE of INRAE (PhenoArch, Montpellier, FR; Cabrera-Bosquet *et al*., 2016). Final leaf number was measured under well-watered conditions in a field environment in Puerto Vallarta, MX. Both experiments were performed under SD conditions, where daylengths throughout the growth cycle never exceeded 12.5 h. Further details for these experiments and the estimation of genotypic effects can be found in Supplemental Information (Methods S2).

(D3) This dataset contains the geographical coordinates and planting dates for 671 field environments across the Northern Hemisphere (latitudinal range of 13.7578 °N to 54.2900 °N) assembled from previous and new projects (File S3).

(D4) This dataset contains a collection of seven temperate inbred genotypes (B37, B73, M37W, Mo17, Oh43, LH123Ht, and 2369) and seven tropical inbred genotypes (CML10, CML258, CML277, CML341, CML373, Tzi8, and Tzi9) measured for days to anthesis in 37 field experiments (File S4). Assembled from previous and new projects, it represents a multi-environment trial (MET) dataset designed to maximize the presence of these 14 genotypes, which were also included in dataset D2. However, their representation across environments was uneven, with temperate genotypes tested in 15 to 22 environments and tropical genotypes in 18 to 37 environments.

(D5) This dataset contains a collection of 109 temperate, 66 tropical, and 61 admixed inbred genotypes, representing separate breeding pools (panel B; Liu *et al*., (2003); Flint-Garcia *et al*., (2005)), measured for days to anthesis across an MET with a maximum of 19 field experiments (File S5). It was assembled from previous projects with an imbalanced representation of photoperiod conditions, including a maximum of three short-day (SD) environments (maximum photoperiod <12.4 h) and 16 long-day (LD) environments (minimum photoperiod >14.5 h), spanning latitudes from 18.00 °N to 42.76 °N.

### Environmental data

For field environments considered in this study, temperature data were acquired from ground and satellite data sources for a common reference point of 2 m height. Ground data were obtained mostly from weather stations at a given field location. In two cases, ground data from a nearby airport within 5 km of the experimental field site was obtained from Weather Underground (https://www.wunderground.com/). Satellite data (weather variable “T2M”; spatial resolution of ½° latitude x ⅝° longitude) was acquired from the R package *nasapower* (Sparks, 2018) using the latitude and longitude coordinates for each field location.

The photoperiod based on civil twilight at a given location for a given day of the year was acquired from the R package *TrenchR* (Buckley *et al*., 2023). This uses the CBM daylength model described by Forsythe *et al*. (1995).

### Comparison of temperature response functions

Throughout this study, calendar days were converted into thermal time to account for growth and development variations caused by temperature differences across field environments. We evaluated three temperature response functions for calculating thermal time using Dataset D1 (Methods S1, Eq. S1-S3). This comparison was based on the assumption that temperature-driven growth and development primarily determine the time to flowering in photoperiod-insensitive genotypes. Under this assumption, temperature differences alone should explain most of the variation in days to anthesis across environments. The function that produces the most consistent thermal time to anthesis values for a given genotype across environments would, therefore, be considered the best. Dataset D1 was used for this assessment since it consisted of temperate-adapted maize hybrids (generally known to be photoperiod insensitive) tested across many independent field environments.

For each of the 24 hybrids, BLUEs for days to anthesis from each field experiment were converted into thermal time using each function. Then, the normalized root mean square error (NRMSE, normalized by the difference between maximum and minimum values) of function-specific thermal times was computed for each genotype. The results were compared to identify the function that minimized the NRMSEs overall. For the purpose of comparing different models for thermal time estimation with the same dataset, we assumed the influence of any latent variables on growth and development could be ignored.

### Definition of a physiological reaction norm for flowering time

The PRN-FTP was reformulated from a previous model by Kiniry *et al*. (1983*a*) that described the relationship between photoperiod (x-axis) and thermal time from crop emergence to tassel initiation 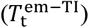 based on temporal dissections of shoot apical meristems (y-axis). Kiniry *et al*. (1983*a*) used growth chambers in which constant levels of photoperiod were applied throughout the plant”s growth cycle, so *DL*^s^ had a known value in each treatment. However, in field environments photoperiods are not constant (*DL*^s^ depends on the day of year within a given environment when a plant become sensitized to photoperiod) and estimating tassel initiation from dissections of meristems is not practical at large scale (rather, in METs, days to flowering are routinely recorded). Besides, as mentioned above, even direct observation of tassel initiation does not correspond to the timing when photoperiod sensitive genotypes become sensitized to photoperiod in LD environments. Therefore, we devised an approach for predicting this unobservable stage of photoperiod sensitization, such that *DL*^s^ could be determined for a given genotype in a given environment. Thus, adapted for METs that cover SD and LD environments, this study defined the PRN-FTP as the relationship between a new proxy measure for *DL*^s^ (x-axis) and thermal time from crop emergence to anthesis (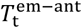;y-axis).

#### Procedure for DL^s^ *Estimation*

There are two main steps used for determining *DL*^s^. First, the 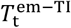 is estimated for genotypes grown under SD conditions in which a photoperiod response is not triggered. In this study, an ecophysiological formalism was used to predict 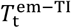 (Eq. 1) from experimental data. This is expected to closely approximate the amount of thermal time it takes for a given genotype to become sensitized to photoperiod, which has been referred to as the basic vegetative phase (BVP; Kiniry *et al*., 1983*b*). Second, the calendar day on which the total thermal time for the BVP is reached is determined using temperature records for a given environment. The daylength for that specific day is then obtained for *DL*^s^. As applied in this study, the two steps of this procedure are as follows:

Step 1: An ecophysiological formalism (Eq. 1) was used to estimate BVP for the 284 genotypes in panel A that were phenotyped in SD experiments (Dataset D2; Methods S2). Therefore, even though the panel included photoperiod sensitive genotypes, their photoperiod responses were not triggered, such that estimates of 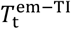 correspond to the BVP. Considering that tassel initiation occurs on the day when the last leaf primordium is formed (Lejeune and Bernier, 1996) and that rates of leaf primordia formation and leaf tip appearance are proportional (Padilla and Otegui, 2005), 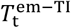 can be calculated as:

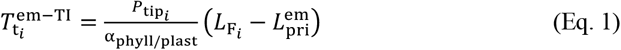

where, 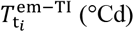 is the cumulative thermal time from crop emergence to tassel initiation of the *i*^th^ genotype, 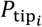 (leaf °Cd^1^) is the phyllochron of the *i* genotype (i.e., the thermal time between the appearance of two successive leaf tips), 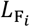 (leaf) is the final leaf number of the *i* genotype, α_plast/phyll_ (dimensionless) is a constant ratio of phyllochron to plastochron (i.e., the thermal time between the formation of two successive leaf primordia), and 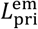 (primordium) is a constant number of primordia formed at crop emergence. Genotype-specific values for 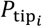 and 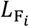 were estimated for panel A (Dataset D2; Methods S2). For maize, α_plast/phyll_ may range from 1.58 to 1.85 (Kiniry, 1991; Lejeune and Bernier, 1996; Padilla and Otegui, 2005; Andrieu *et al*., 2006). In this study, we set it at the average value reported in the literature (1.73). There are four to five leaf primordia in the embryo of maize seeds (Bonnett, 1954; Padilla and Otegui, 2005), and between germination and crop emergence about two other leaf primordia are formed (Padilla and Otegui, 2005). Thus, here, 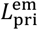 was set equal to 5.5 primordia.

Step 2: Using 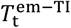 for the BVP of a given genotype, along with emergence date and temperature records from a given environment, the calendar day on which the genotype becomes sensitized to photoperiod was predicted. When observed emergence dates were unavailable, a thermal time of 76 °Cd was assumed for the period from sowing to emergence. This was estimated from observed data for 15 environments within Dataset D4 (sourced from the Maize ATLAS project; see Table S1), calculated using the temperature response function Eq. S2.

#### Thermal time from crop emergence to anthesis (y-axis of the PRN-FTP)

For each genotype in each environment, 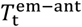 was calculated as the cumulative thermal time between the dates of emergence date (observed or predicted) and anthesis (determined from BLUEs of days to anthesis).

### Envirotyping sensed photoperiod

Dataset D3 was used to computationally envirotype *DL*^s^ across 671 environments in the Northern Hemisphere. In each environment, *DL*^s^ was envirotyped for three maturity levels defined by the lower (5^th^; early maturity), median (50^th^), and upper (95^th^; late maturity) percentiles of the BVP distribution estimated for panel A (Dataset D2; Table 1). The difference in *DL*^s^ between maturity levels was calculated to assess how genotypes with different BVPs may vary in the photoperiod they perceive within a given environment. Knowing that daily changes in photoperiod increase with latitude, the per-environment differences in *DL*^s^ were compared to rates of photoperiodic change. The rate of photoperiodic change within each environment was calculated from photoperiods between the days immediately before and immediately after *DL*^s^ for the 50^th^-percentile maturity level.

**Table 1.** Percentiles of the distribution for the basic vegetative phase (BVP) in panel A.

|  | **Min** | **5%** | **25%** | **50%** | **75%** | **95%** | **Max** |
| --- | --- | --- | --- | --- | --- | --- | --- |
| **BVP (°Cd)** | 169 | 239 | 286 | 326 | 375 | 434 | 496 |

When modeling the PRN-FTP for each genotype from MET data, 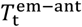 (response variable) values will correspond to the environment-specific observations for each genotype, while *DL*^s^ (predictor variable) could either be based on the BVP for the same particular genotype or a common reference representing all genotypes. The choice depends first on whether BVP data are available for all genotypes in the MET, but also whether different genotypes in the same environment experience large or small differences in photoperiod.

### Statistical analysis of the PRN-FTP

Two different regression functions, bilinear and linear, were used for modeling the PRN-FTP. This involved two different approaches as follows:

#### Bilinear function for estimating three parameters of the PRN-FTP

With a sufficient density of data and coverage of *DL*^s^ spanning SD and LD environments, photoperiod sensitive genotypes are expected to follow a bilinear pattern of response. In SD environments 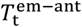 remains constant, and above a critical threshold for LD environments it increases linearly with photoperiod. For this, the PRN-FTP can be modeled as a continuous two-phase (bilinear) regression function with threshold detection (Fong *et al*., 2017*a*). We used the “hinge model” in the R package *chngpt* (Fong *et al*., 2017*b*), defined as follows:

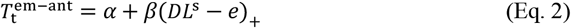

where 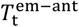 (described before) is the response variable; *e* is the threshold parameter; *DL*^s^ is the sensed daylength with a threshold effect; *I*(*DL*^s^ > *e*) = 1 when *DL*^s^ > *e* and 0 if *DL*^s^ ≤ *e*; (*DL*^s^ − *e*)_+_ denotes the hinge function, which equals *DL*^s^ − *e* for *DL*^s^ > *e* and 0 otherwise; and α and β are the intercept and slope parameters, respectively. The model was fitted using default settings for *chngpt*, with no constraints set for estimating the threshold value (breaking point of the bilinear model). Estimated effects include an intercept, threshold value, and slope, which correspond to physiological components of flowering time: flowering time *per se*, a critical threshold for photoperiod sensitivity, and photoperiod sensitivity, respectively.

#### Linear function for estimating two parameters of the PRN-FTP

When fitting the bilinear model is not feasible due to insufficient density of data, inadequate coverage of *DL*^s^ for threshold detection, or because it is not an appropriate model for all genotypes (i.e., photoperiod insensitive genotypes), we modeled the PRN-FTP as a linear regression function, forgoing threshold estimation. So that the intercept for 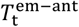 is estimated at the mean *DL*^s^ of SD environments rather than the shortest day environment, environments across the MET were first classified as SD or LD based on whether *DL*^s^ falls below or above a fixed threshold, respectively (we used 13.5 h which corresponded to the average threshold estimated from the analysis of Dataset D4, see results). The predictor values for each environment were then recoded as the mean *DL*^s^ value for the respective SD and LD environments in the regression model. Finally, when specifying the linear regression model, the SD and LD *DL*^s^ means are offset with the mean SD-*DL*^s^ value set to zero to obtain the proper intercept estimate for the corresponding genotype. Hereafter, we refer to this as the “LD-SD” method for modeling the PRN-FTP. This is similar to prior studies that used the difference between mean thermal times to flowering in LD and SD environments as an estimate of photoperiod senstivity (Coles *et al*., 2010; Hung *et al*., 2012), except here an intercept for flowering time *per se* (in units °C) and slope for photoperiod sensitivity (in units °Ch^-1^) are estimated.

#### N.B

Two particular points are relevant for interpreting parameters estimated by the LD-SD method. First, envirotyping of *DL*^s^ to obtain the mean values for SD and LD environments may be based on each specific genotype being modeled or a common reference genotype (this point is described in other parts of the study). Even if a common reference BVP is used to envirotype *DL*^s^ for all genotypes, the mean values for *DL*^s^ can vary due to imbalanced observations of the genotypes across environments, which occurs frequently in MET data. That is, although *DL*^s^ may be envirotyped for a common reference BVP in each environment, the regression analysis is performed using predictor values linked to the observed 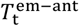 data being modeled for each genotype. Second, for photoperiod sensitive genotypes, it is expected that estimates of photoperiod sensitivity will be biased as a function of the deviation between the SD-*DL*^s^ value and the true (unknown) critical photoperiod for a given genotype. When the true critical photoperiod is less than the SD-*DL*^s^ value, the estimate will be upward biased, and when it is greater it will be downward biased.swart

#### Correction for latent variables affecting thermal times to anthesis

If only temperature influenced the flowering time of a photoperiod insensitive genotype, we expect its 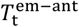 would be nearly identical across all environments in an MET. However, aside from photoperiod sensitivity, misrepresentation in the temperature response function, or measurement error, deviations in 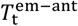 could be due to various possibilities that vary from one environment to another, including the influence of other environmental factors, differences in crop management practices, differences between air temperature and the temperature sensed by the crop, imprecision and biases in weather data, or other latent variables. Therefore, with dataset D4, we tested an alternative approach using photoperiod insensitive control genotypes to adjust for the effects of latent variables across environments prior to constructing the PRN-FTP.

Seven photoperiod insensitive genotypes were defined as control genotypes in Dataset D4. The 37 environments comprising the MET were assembled from different experiments. Unfortunately, all seven control genotypes were not present in all environments, so two sets of control groups were defined (set 1: B37, B73, M37W, Mo17, and Oh43; set 2: LH123Ht and 2369). Sets 1 and 2 were associated with 22 and 15 of the environments, respectively. For each set, the overall mean of 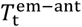 was calculated across environments of the MET in which they were present. The environment-specific residuals were then used to adjust the thermal times for the remaining non-control genotypes in each corresponding environment. We note that using two sets of control genotypes that were also imbalanced across environments (see the earlier description of Dataset D4) is not ideal and could introduce some bias, but any improvement in the model fit would presumably reflect positively on this approach. Therefore, using Dataset D4, we compared the overall RMSE of the PRN-FTP model fit and standard errors of parameter estimates before and after applying the correction method.

### Applications of the PRN-FTP

Datasets D4 and D5 were used to test the concepts developed in this study for modeling and analyzing the PRN-FTP from MET data.

#### Methodological evaluation

The MET for Dataset D4 had a sufficient coverage of *DL*^s^ for fitting the bilinear model (assumed to be the biologically correct one). Additionally, BVP values from Dataset D2 could be used to obtain genotype-specific *DL*^s^ values for the genotypes in D4. This allowed us to test different approaches to modeling the PRN-FTP, including: (i) the impact of using latent-variable adjusted values of 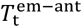;(ii) the impact of envirotyping *DL*^s^ using genotype-specific values versus a common reference value for the BVP; and (iii) the impact of using the bilinear model versus the LD-SD method on the estimated parameters for flowering time *per se* and photoperiod sensitivity (the threshold for critical photoperiod can only be estimated with the bilinear model). Although this was performed with only seven tropical maize genotypes, analysis with the bilinear model also provided a preliminary survey of potential variation in the three PRN-FTP parameters for maize.

#### Analysis of maize diversity

We sought to investigate the wider variation in PRN-FTP parameters in a diverse collection of 214 maize genotypes adapted to different geographical zones (Dataset D5). However, due to both an insufficient density and coverage of *DL*^s^ and the mixed composition of photoperiod sensitive and insensitive genotypes, the bilinear function could not be fit. This was true even for genotypes known to be photoperiod sensitive, such as those used for methodological evaluation (we observed bogus estimates of the parameters). Moreover, most of the genotypes were not present in Dataset D2 so genotype-specific BVP values for *DL*^s^ envirotyping could not be systematically used. Therefore, for Dataset D5, the LD-SD method was applied, with *DL*^s^ values obtained using the BVP for the 50^th^-percentile reference maturity determined with Dataset D2 (Table 1). The regression model was fit using latent-variable adjusted BLUEs 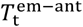 regressed onto mean *DL*^s^ for the SD and LD environments as described in the previous section. Latent variable adjustment was performed using the set 1 control group (indicated above) common across the Dataset D5 MET.

## Results

### Choosing a temperature response function to construct the PRN-FTP

Thermal time is used to determine *DL*^s^ for envirotyping along with 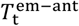 for modeling the PRN-FTP, such that different temperature response functions for calculating thermal time could result in different outcomes. A comparison of three different temperature response functions showed no significant difference in the NRMSEs for thermal times to anthesis for 24 photoperiod-insensitive hybrids tested in 25 to 26 environments in North America. Regardless of the temperature response function that was tested, for each hybrid, the environment-specific thermal time to anthesis deviated, on average, by ∼10% (range 9 to 13%) from the across-environment mean (Fig. S1). Therefore, for the current study, we chose the non-linear response function by Wang and Engel (1998) adapted for maize (Eq. S2) due to its common implementation in crop growth models (Wang *et al*., 2017), its higher physiological realism compared with the linear response function (Eq. S1), and its greater flexibility compared to the Arhenus model (Parent *et al*., 2010).

### Predicting when maize becomes sensitized to photoperiod

There is no known visible marker to measure the timing when maize plants become sensitized to photoperiod for determining *DL*^s^. Therefore, we defined this using the BVP (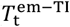 in SD conditions), which closely corresponds to the thermal time required for plants to reach their photoperiod-sensitized stage (Kiniry *et al*., 1983*b*; Bassiri *et al*., 1992). An issue is that directly observing tassel initiation requires temporal dissections of shoot apical meristems which is not possible at a large scale. To circumvent this, we used an ecophysiological formalism to estimate 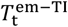 (Eq. 1) for the 284 genotypes in panel A grown under SD conditions (Dataset D2; File S2). A separate SD experiment validated this for eight of the genotypes using microscopy data on differentiation of shoot apical meristems into tassel primordia (Methods S2). The Pearson correlation coefficient between predicted and observed BVP values was 0.84, indicating the formalism-based predictions for 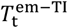 align closely with observational data. Table 1 summarizes percentiles of the BVP distribution for panel A, with the 5^th^, 50^th^, and 95^th^ percentile values used to represent early, medium, and late maturity groups for envirotyping *DL*^s^.

### Spatiotemporal variation in the sensed photoperiod for maize

Envirotyping with *DL*^s^ can be used to understand how different genotypes sense photoperiod across and within environments, and how this varies according to spatiotemporal patterns in climate and weather. To examine how photoperiods are perceived by maize, we computationally envirotyped *DL*^s^ for 671 field environments across the Northern Hemisphere where maize has been previously grown (Fig. 2).

**Figure 2.**
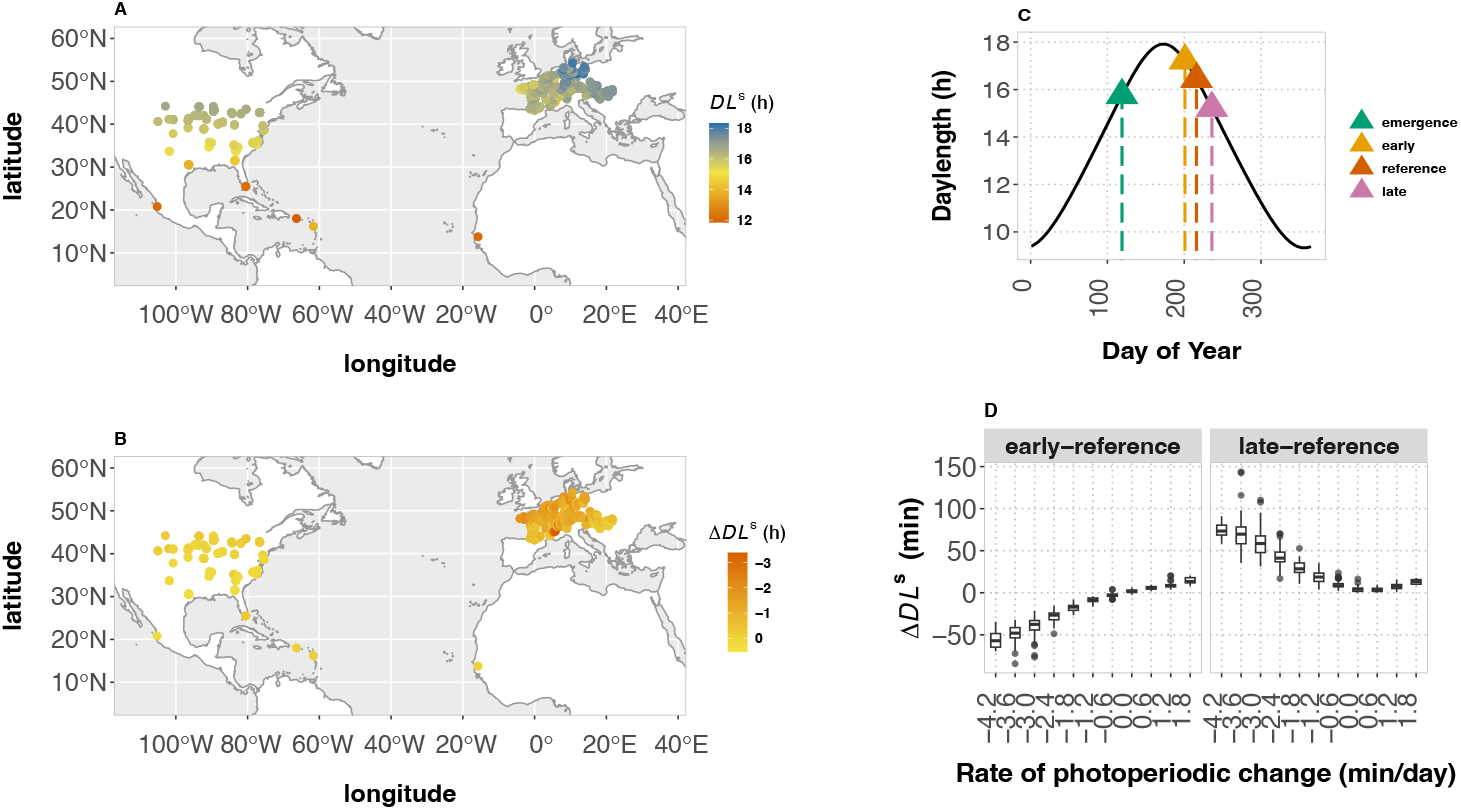
Envirotyping of sensed photoperiod across and within maize fields in the Northern Hemisphere. (A) Sensed photoperiod (*DL*^s^ for the 50^th^-percentile maturity reference) across maize field sites in the Northern Hemisphere. (B) Within environment differences in hours of sensed daylength (Δ*DL*^s^) between the 95^th^ and 5^th^ percentile maturity groups defined by the BVP distribution of panel A. (C) The figure portrays how different maturity groups can reach the photoperiod sensitized stage on different days of the year within the same environment, leading to differences in *DL*^s^ (the 5^th^, 50^th^, and 95^th^ percentile values of the BVP in panel A are indicated as early, reference, and late maturity groups) (D) The relationship between within environment differences in *DL*^s^ and rates of daily photoperiodic change. Here, the differences are shown in minutes for comparisons between the reference maturity group relative to the early and late maturity groups. As reflected in (B), the boxplots show that genotypes within the same environment experience increasingly greater differences in photoperiods when rates of photoperiodic change increase, which occurs in association with increasing latitude.

We first summarized the spatial axis of variation in *DL*^s^ using the 50^th^ percentile of 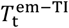 (326 °Cd) as a common reference for envirotyping (Table 1; Fig. S2). The range in *DL*^s^ across the entire set of environments evaluated was 11.9 to 18.3 h, a difference of 6.4 hours (Fig. 2a). We compared the *DL*^s^ for maize above 30° N in the USA and Europe, corresponding to regions where maize is used extensively for scientific research and agricultural production (Figure 2a-b). Mean *DL*^s^ across the USA was 16.1 h and in Europe was 16.7 h, with lower (5-percentile) to upper (95-percentile) ranges of 14.3 to 16.4 and 15.7 to 17.5, respectively. These spatial patterns are primarily driven by climatic variation in photoperiod across latitude coupled with the seasonality for maize cultivation. Since temperature can speed or slow growth and development, interannual differences in temperature and planting dates at the same location can give rise to additional variation in *DL*^s^ for maize. To examine this, we selected 118 field sites where maize was planted for at least two years and found that *DL*^s^ within a single location varied by as much as 1.7 hours, with a mean absolute difference of 0.3 h.

Next, we questioned the extent to which multiple genotypes grown within the same field environment perceive a different photoperiod. Beyond the equator, photoperiod is not constant across days of the year, with increasing rates of daily change in association with latitude. Therefore, genotypes differing in BVP that reach their photoperiod-sensitized stage on different days would perceive a different photoperiod (Fig. 2b-c), but to our knowledge the magnitude by which this varies has not been investigated. Using the 5^th^ (239 °Cd) and 95^th^ (434 °Cd) percentiles of the BVP distribution (Table 1) to represent early and late percentile-maturity groups, the *DL*^s^ at these limits frequently differed by more than 1 h, with the largest difference reaching 3.8 h (Fig. 2b). Differences in *DL*^s^ above 2 h were rare and occurred only in locations at high latitudes where late genotypes are not cultivated for maize production. Nevertheless, this highlights how the genetic diversity of maize can intersect with environmental variation within the same field, leading to differences in the photoperiod to which genotypes respond.

If the 50^th^-percentile for BVP is used as a common reference for envirotyping *DL*^s^, the magnitude of inaccuracy compared to genotype-specific *DL*^s^ varies according to the specific set of genotypes used and the environment in which they are tested. For instance, the early and late percentile-maturity groups differed from the 50^th^-percentile reference by as little as no difference to as much as 1.4 h and 2.4 h, respectively (File S3). The larger differences occurred at higher latitudes where daily rates of photoperiodic change were greatest. As expected, this difference reduced to zero as a function of decreasing rates in photoperiodic change (Fig. 2d).

### Using METs for modeling the PRN-FTP

To test approaches for modeling the PRN-FTP from METs, we combined flowering time data from multiple projects totaling 37 field experiments (Dataset D4) that included, albeit with imbalanced data (Table S1), a common set of seven genotypes of tropical origin that were expected to be photoperiod sensitive. This was used to compare procedures for estimating genotype-specific PRN-FTP parameters for flowering time *per se* (intercept), critical photoperiod (threshold), and photoperiod sensitivity (slope) using a two-phase regression model (see Methods; Fig. 3).

**Figure 3.**
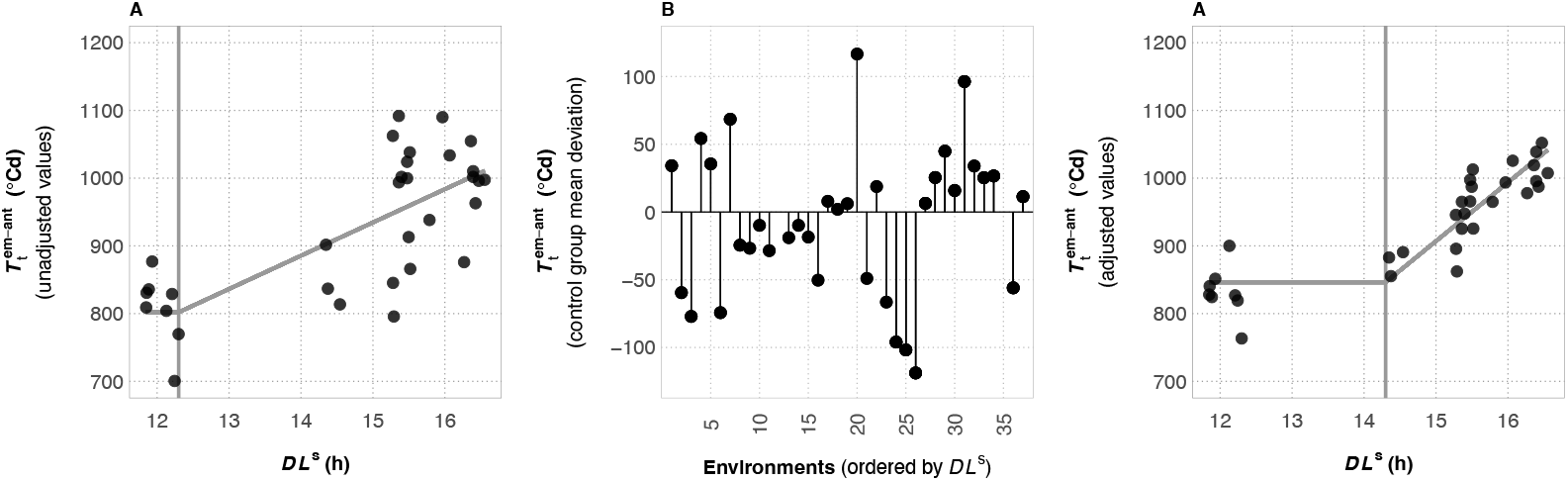
Adjusting for latent variable effects across a multi-environment trial network when fitting the PRN-FTP. (A) The plot shows the PRN-FTP for genotype CML341, using unadjusted BLUEs for thermal time from crop emergence to anthesis (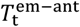; black points). The grey lines show the estimated flowering time *per se* (horizontal line), critical threshold value (vertical line), and photoperiod sensitivity (sloped line). (B) Deviations from the overall average 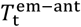 for the control groups of photoperiod insensitive genotypes estimated across the same environments, with the environment identifiers ordered as in plots (A) and (B) according to sensed daylength (*DL*^s^). (C) The PRN-FTP for genotype CML341 after adjusting the BLUEs by deviations in (B), leading to different estimates for the flowering time *per se*, the critical threshold value, and the photoperiod sensitivity.

Using MET data to construct the PRN-FTP makes the simplistic assumption that physiological sensitivities to temperature and photoperiod are the only environmental factors influencing flowering time. Although these are dominant factors, other variables can also affect maize growth and development, but the specific causes are often unknown and likely to vary from one field environment to another. Therefore, we devised an approach to control for latent effects on flowering time and evaluated how adjusted values for the 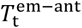-phenotypic axis affected PRN-FTP parameter estimation. Using a designated set of photoperiod-insensitive control genotypes present in the D4 MET, their overall mean 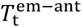 across environments was defined as an expected value for the specific effect of temperature on flowering time (i.e., for photoperiod insensitive genotypes, 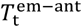 is expected to be the same across environments, with deviations attributed to latent variables; see Materials and Methods). Then, environment-specific deviations from the overall mean were used to adjust 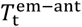 values for the remaining genotypes in each of the corresponding environments. Fig. 3 portrays this procedure and how it can affect estimates of the PRN-FTP parameters.

When tested using genotype-specific *DL*^s^ values for each of the seven tropical genotypes, latent variable adjustment of 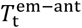 enhanced the precision of the overall model fit of the PRN-FTP by an average of two-fold (Table 2). Using adjusted values also led to changes in the estimated thresholds of sensitivity, which in turn resulted in differences in the estimated flowering times *per se* by as much as 5% and slopes of sensitivity by as much as 77% (Table 2).

**Table 2.** Impact of latent variable adjustment and genotype-specific envirotyping on PRN-FTP parameter estimation for different maize genotypes.

| Genotype | Intercept<br>(°Cd) | Threshold<br>(h) | Slope<br>(°Cd h<sup>-1</sup>) | RMSE<sup>a</sup> |
| --- | --- | --- | --- | --- |
| **Unadjusted values with genotype-specific *DL*<sup>s</sup>** |  |  |  |  |
| CML10 | 851.0 ± 28.6 | 12.3 ± 0.6 | 44.3 ± 9.7 | 82.0 |
| CML258 | 934.6 ± 33.4 | 12.3 ± 0.7 | 42.1 ± 11.4 | 95.4 |
| CML277 | 856.5 ± 22.1 | 14.5 ± 0.6 | 96.7 ± 18.7 | 76.0 |
| CML341 | 802.0 ± 24.7 | 12.3 ± 0.6 | 49.1 ± 8.4 | 71.2 |
| CML373 | 815.2 ± 30.6 | 12.3 ± 1.1 | 35.9 ± 10.7 | 67.5 |
| Tzi8 | 858.0 ± 29.1 | 12.4 ± 1.1 | 30.9 ± 9.9 | 86.4 |
| Tzi9 | 809.0 ± 29.5 | 12.2 ± 1.1 | 26.3 ± 9.3 | 85.0 |
| **Adjusted values with genotype-specific *DL*<sup>s</sup>** |  |  |  |  |
| CML10 | 872.8 ± 16.7 | 12.3 ± 0.6 | 36.9 ± 5.6 | 47.7 |
| CML258 | 953.8 ± 21.2 | 12.3 ± 0.9 | 37.7 ± 7.2 | 60.5 |
| CML277 | 869.8 ± 9.2 | 14.5 ± 0.6 | 87.7 ± 8.0 | 31.3 |
| CML341 | 846.0 ± 9.8 | 14.3 ± 0.6 | 86.7 ± 7.9 | 33.8 |
| CML373 | 853.1 ± 11.6 | 12.3 ± 0.8 | 26.4 ± 4.0 | 25.6 |
| Tzi8 | 888.8 ± 14.7 | 14.3 ± 1.0 | 44.3 ± 10.7 | 52.8 |
| Tzi9 | 842.1 ± 11.2 | 14.4 ± 0.9 | 41.8 ± 8.8 | 40.0 |
| **Adjusted values with common reference *DL*<sup>s</sup>** |  |  |  |  |
| CML10 | 873.7 ± 16.5 | 12.3 ± 0.6 | 37.5 ± 5.7 | 47.6 |
| CML258 | 955.8 ± 21.0 | 12.3 ± 0.9 | 37.7 ± 7.3 | 60.5 |
| CML277 | 870.2 ± 9.3 | 14.3 ± 0.5 | 77.0 ± 7.1 | 32.1 |
| CML341 | 846.9 ± 9.9 | 14.3 ± 0.6 | 83.2 ± 7.6 | 34.2 |
| CML373 | 853.5 ± 11.5 | 12.2 ± 0.8 | 26.3 ± 4.0 | 25.5 |
| Tzi8 | 888.6 ± 14.7 | 14.5 ± 1.1 | 47.9 ± 11.6 | 52.8 |
| Tzi9 | 841.9 ± 11.2 | 14.5 ± 0.8 | 43.2 ± 9.0 | 39.9 |

<sup>a</sup>Root mean squared error (RMSE) is reported for the full bilinear model.

As described in the previous section, genotypes with a different BVP can experience different photoperiods within environments, such that points along the *DL*^s^-environmental axis of the PRN-FTP may differ among genotypes in a common MET. This caused us to question the standard approach to modeling reaction norms from METs where all genotypes are assumed to experience the same environmental conditions in each field (i.e., values for a given environmental covariate is assumed to be common for all genotypes). If the specific conditions experienced by different genotypes are sufficiently large, this could impact parameter estimates of a reaction norm. To test this, we compared modeling the PRN-FTP using *DL*^s^ obtained from the BVP for each genotype versus the 50^th^-percentile BVP as a common reference. The seven genotypes presented adequate variation for testing this: compared to the 50^th^-percentile reference point, genotype-specific values for the BVP ranged from 266 ºCd to 399 ºCd, corresponding to the 16^th^ and 89^th^ percentiles of the BVP distribution estimated from 284 genotypes (Table 1). However, the set of environments in the MET network of Dataset D4 presented smaller contrasts in *DL*^s^ than would occur at higher latitude (European) environments (Fig. 2b): the largest difference in *DL*^s^ between these specific genotypes and the common reference was 0.22 h. In this case, changes in coefficient estimates for thresholds of sensitivity and flowering time *per se* were negligible, but the slope parameter for photoperiod sensitivity changed by 12% for one of the gentoypes (Table 2). Nevertheless, these results show that intra-environment variation in *DL*^s^ among genotypes can affect PRN-FTP parameters estimated from MET data. Even for this small sample of genotypes, Fig. 4 highlights a wide range of diversity for flowering time plasticity in maize consisting of unique parameterizations of the PRN-FTP.

**Figure 4.**
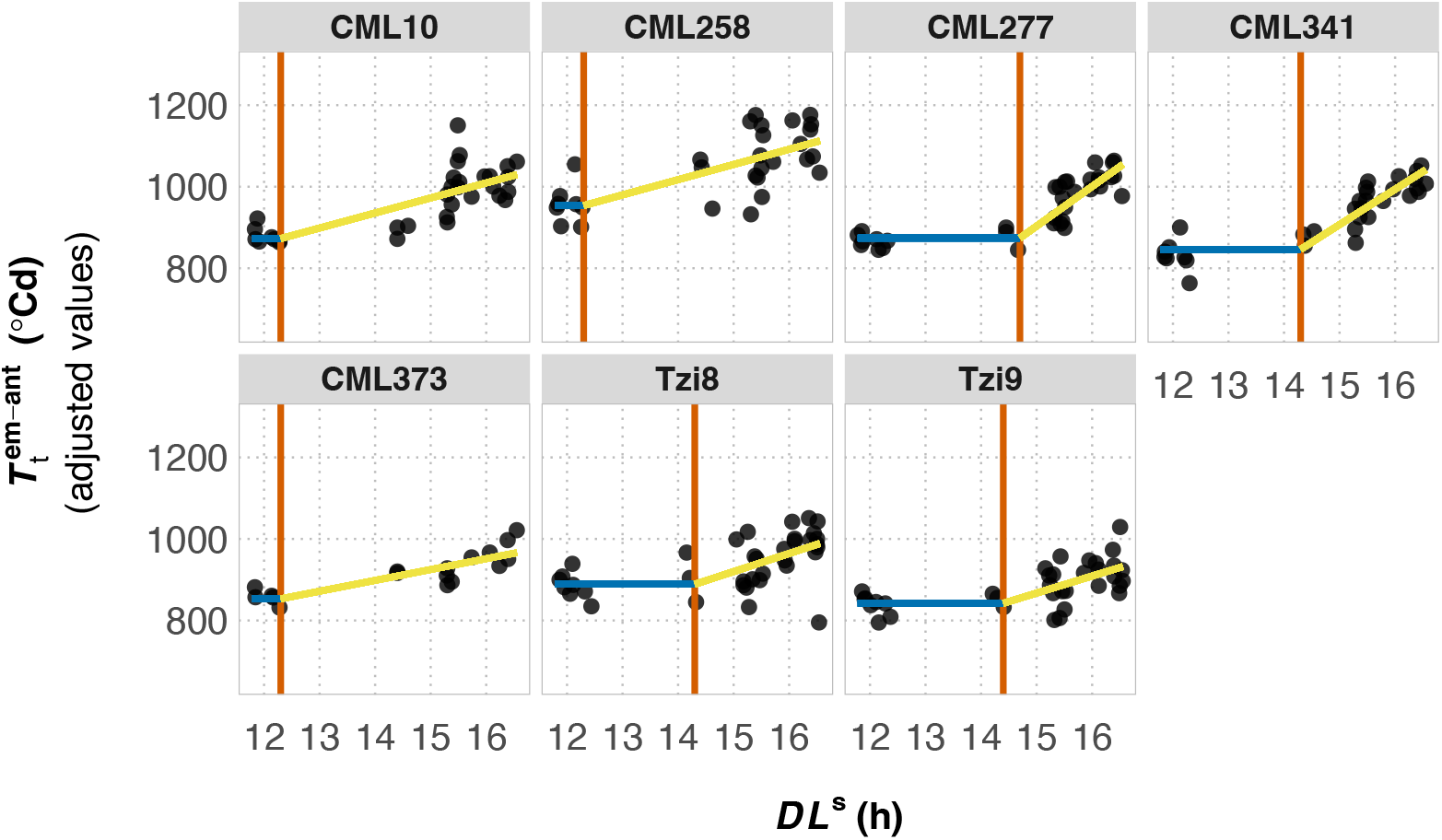
The PRN-FTP for seven tropical genotypes of maize. (A) For each genotype, the plots show the three PRN-FTP parameters estimated from the bilinear function fitted using latent-variable adjusted BLUEs for thermal time from crop emergence to anthesis 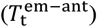 in relation to sensed daylength (*DL*^s^). Parameter estimates are shown for the flowering time *per se* (blue line), the critical threshold value (red line), and the photoperiod sensitivity (yellow line).

### Relating components of the PRN-FTP to understand maize adaptation

Since flowering time *per se* and photoperiod sensitivity underlie flowering time adaptation across environments (Choquette *et al*., 2023), we questioned how the trait space of these PRN-FTP parameters is structured among primary breeding pools of maize. For this, we assembled an MET dataset (D5) from 19 field trials with a common set of genotypes consisting of tropical, temperate, and admixed breeding material. Computational envirotyping showed that *DL*^s^ for the D5 MET was restricted to two small intervals of photoperiod conditions, with three environments of 11.9 to 12.4 h photoperiod and 16 environments of 14.5 to 16.5 h photoperiod. With this density of data and gap in coverage of *DL*^s^, the bilinear function for the PRN-FTP could not be reliably fit. Besides, the collection of material included photoperiod insensitive genotypes that do not follow a bilinear response to photoperiod. In this case, the PRN-FTP was modeled using an LD-SD method based on linear regression after, as previously described, adjusting for latent variables with a designated set of control genotypes. Examining the relationship between flowering time *per se* and photoperiod sensitivity showed that temperate and tropical material occupy distinct territories of the trait space (Fig. 5; File S6). Temperate genotypes tend to have a lower flowering time *per se* and nearly no photoperiod sensitivity, while tropical genotypes tend to have a higher flowering time *per se* coupled with a large range in photoperiod sensitivity. Fitting genetic expectations for polygenic traits, genotypes with admixed ancestry are positioned between these two regions of the trait space.

**Figure 5.**
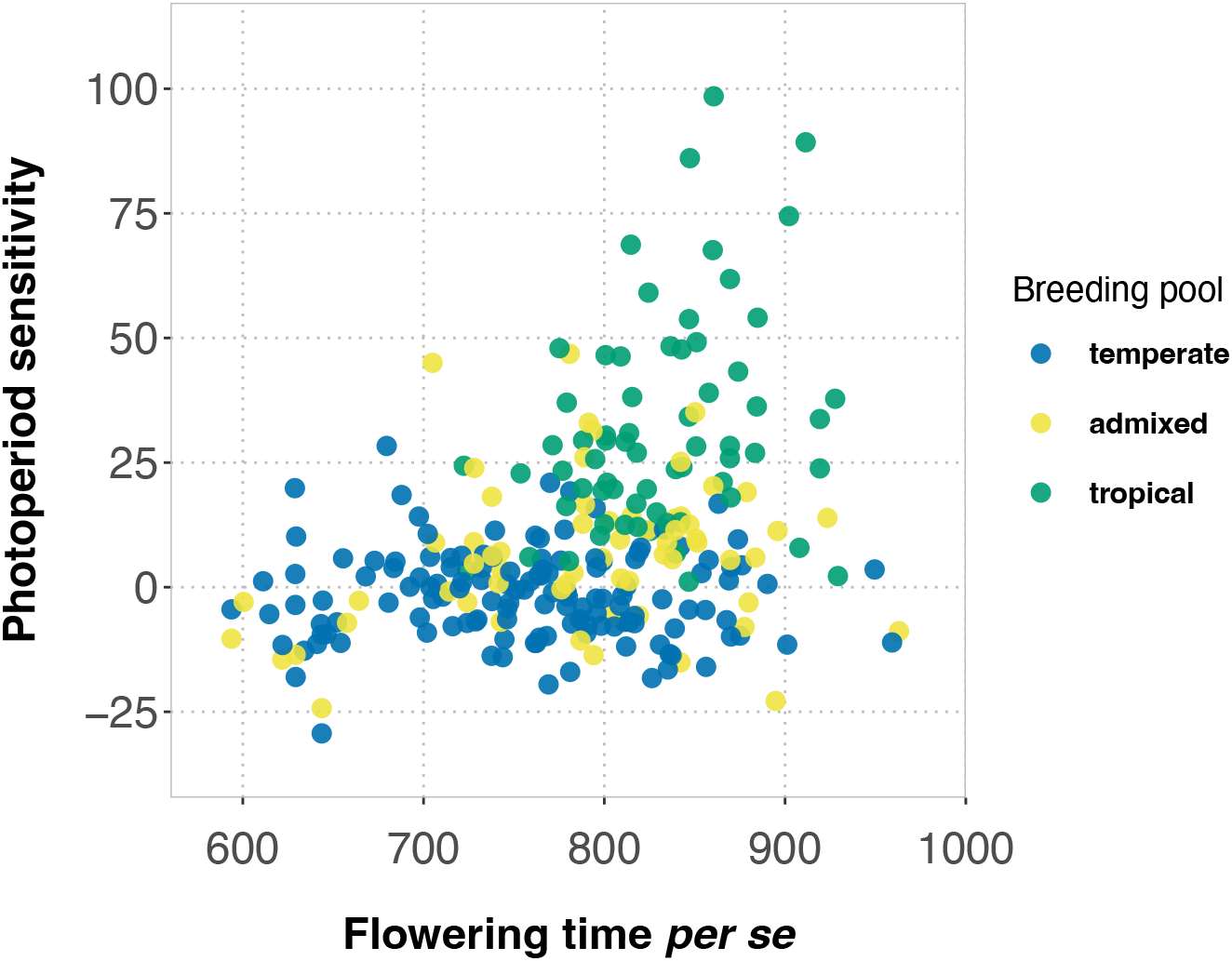
The trait space for flowering time *per se* and photoperiod sensitivity in maize breeding material. Points correspond to genotypic values for flowering time *per se* (ºCd) and photoperiod sensitivity (ºCd h^-1^), estimated using the LD-SD regression method on latent-variable adjusted BLUEs. Breeding pools are represented by genotypes previously assigned to temperate (*n*=137), admixed (*n*=62), and tropical (*n*=66) sources based on structure analysis of marker data (Flint-Garcia *et al*., 2005).

## Discussion

Understanding trait plasticity is important for adapting crops to new environments. To study the variation in plastic responses, METs allow large collections of genotypes to be characterized across environmental gradients. When modeling genotype-environment effects, the assumption—implicit in standard models—is that all genotypes experience the same conditions in each environment (an exception: (Millet *et al*., 2019)). However, as demonstrated in this study, due to phenological variation and stage-specific sensitivities, different genotypes within each field environment may not experience identical environmental conditions. This can complicate the definition of environmental indices for modeling reaction norms from METs, challenging standard analyses that assume common environmental conditions are sensed by all genotypes. Based on the ecophysiology of maize flowering time, the current study defines a new metric, *DL*^s^, for quantifying the specific photoperiods sensed by different genotypes within and across environments. Using *DL*^s^ to envirotype maize fields in the Northern Hemisphere, we showed that, at high latitudes, where cooler temperatures that slow development combine with relatively large daily rates of photoperiodic change, genotypes in the same environment can experience more than a 1 h difference in *DL*^s^ (Fig. 2). This occurs more often and is more pronounced across maize environments in Europe than in the U.S. due to higher rates of daily photoperiodic change (Fig. 2d). Thus, to provide more accurate information about the variation in trait plasticity, the construction of genotype-specific reaction norms for flowering time and other traits should be considered on a case-by-case basis.

Constructing the PRN-FTP from METs is the most practical approach for studying the diversity of flowering time plasticity, but this is challenged by the geographical structure of photoperiod (Fig. 2) and the sensitivity of parameter estimation to gaps in *DL*^s^ (Table 2, Fig. 4). Testing across numerous field sites spanning nearly an entire hemisphere is required to capture a sufficient range and density of *DL*^s^ to model genotypic variation in the PRN-FTP. We found that currently available MET datasets for maize do span LD and SD conditions but have limited to no coverage of *DL*^s^ from 12.4 to 14.5 h, the specific range for expected variation in the critical photoperiod for maize (Rood and Major, 1980; Kiniry *et al*., 1983*a*). With scant information on critical photoperiods for maize, and given the dependence of accurate threshold estimation for regression analysis of the PRN-FTP (Table 2), envirotyping with *DL*^s^ could be used to design new METs for future research on flowering time plasticity.

The PRN-FTP provides a parsimonious model for flowering time plasticity, taking into account the dominant effects of temperature and photoperiod. However, in METs, latent variables can affect flowering time, requiring appropriate adjustments. In particular, constructing the PRN-FTP from thermal time estimates can be obscured by the influence of other factors besides temperature on growth and development, such as drought (e.g., Harrison *et al*., 2014; Herrero and Johnson 1981). To address this issue, we applied a between-environment covariate adjustment using photoperiod-insensitive control genotypes expected to have the same thermal time to anthesis across LD and SD environments. This consistently resulted in a more precise fit of the bilinear response function for the PRN-FTP (Table 2; Fig. 3). Nevertheless, it is important to acknowledge that the choice of a control group could introduce some bias, as it may uniquely vary in other ecophysiological responses. Another challenge in modeling the PRN-FTP is that not all genotypes exhibit a clear bilinear response. For example, photoperiod-insensitive genotypes only display differences in flowering time *per se*, rendering the bilinear regression function invalid. In such cases, linear regression using the LD-SD method is the most appropriate way of capturing the relationship between 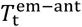 and *DL*^s^. Thus, while the PRN-FTP offers a valuable framework for understanding flowering time plasticity, careful consideration of latent effects, choice of control group, and genotype-specific responses is necessary for accurate modeling. Future work could consider alternative statistics to simultaneously model mixtures of genotypes with bilinear and linear reaction norm structures for the PRN-FTP.

A large body of work has elucidated the GRN-topology of multiple pathways involved in the transduction of signals that control flowering time in plants (Srikanth and Schmid, 2011). Reflective of this complex regulatory system, MET mapping studies have shown the genetic architecture of phenotypic variation in days to flowering to be polygenic (e.g., maize - Buckler et al., 2009). While many genes are involved, using ecophysiology to redefine the context for genetic mapping can clarify the roles of individual loci or genes in relation to the GRN. As one example from maize, mapping variation in the difference between thermal time to flowering in LD and SD environments—an ecophysiological transformation of days to flowering into a measure of photoperiod sensitivity—led to the identification of a causal variant affecting the regulation of the photoperiod sensitivity pathway (Hung *et al*., 2012; Yang *et al*., 2013). The same study by Hung *et al*. (2013) demonstrated that this approach isolates loci specifically responsive to photoperiod from other loci that contribute to variation in flowering time in other ways. Nevertheless, the full topology and context-dependent expression of the GRN regulating flowering time remains unresolved, let alone its connections across physiological to whole-plant scales. We expect that using *DL*^s^-guided MET design to estimate and decompose variation in flowering time *per se*, critical photoperiods, and photoperiod sensitivity will help to further disentangle the genetic architectures of these context-specific traits. By relating these architectures to the GRN, this approach will facilitate testing of hypotheses on the multiscalar relationships and genotype-environnment interactions for flowering time.

Flowering time adaptations have facilitated the evolution and geographical spread of plant species (Gaudinier and Blackman, 2019). The expansion of maize from its tropical origins to shorter growing seasons in the temperate zone was mediated by selection for early flowering time (Vigouroux *et al*., 2008; Tenaillon and Charcosset, 2011; Swarts *et al*., 2017), with evidence of selection on multiple pathways of the GRN (Brandenburg *et al*., 2017). Distinct breeding material was established from locally preadapted populations that have largely persisted as separate pools of tropical and temperate maize (Liu *et al*., 2003). Taken together, we reasoned that the PRN-FTP trait space of flowering time *per se* and photoperiod sensitivity for modern varieties representing these pools would reflect the historical perspective of adaptation and subsequent breeding in maize. Figure 5 corroborates this: assuming tropical genotypes reflect an ancestral state defined by moderate to late flowering time *per se* (adaptation to long growing seasons in the tropics) and segregation for photoperiod sensitivity (weak to no selection in SD environments), temperate maize shows both reduced flowering time *per se* and photoperiod sensitivity (adaptations for a shortened crop cycle). While this general structuring of plasticity between tropical and temperate breeding pools could be expected, Fig. 5 reveals an empty territory of the space where essentially no genotypes exist that are both early *per se* and strongly sensitive to photoperiod. This combination could also condition moderate maturity, for example. We hypothesize that this is a byproduct of the preadaptation history and subsequent breeding of maize, and not indicative of an intrinsic coupling between different pathways of the GRN or physiological controls of flowering time. This is supported by results from experimental evolution for early flowering time in tropical populations under contrasting photoperiod conditions. In LD environments, photoperiod sensitivity and flowering time *per se* are co-selected but show different rates change across generations, with photoperiod sensitivity rapidly eliminated (Teixeira *et al*., 2015; Choquette *et al*., 2023). In contrast, in SD environments, responses to selection can be attributed to flowering time *per se* with no changes observed for photoperiod sensitivity. This highlights how the PRN-FTP can be used for studying crop diversity, and that knowledge about the historical context of adaptation and breeding can aid in interpreting the observed patterns in the PRN-FTP-derived trait space.

## Conclusions

In this study, we explored various aspects of a physiological reaction norm for modeling flowering time plasticity in maize. Envirotyping with *DL*^s^ is a valuable tool for examining the global distribution of photoperiodic sensing in maize and constructing the PRN-FTP from MET data. Notably, the geographical distribution of current public MET networks for maize are concentrated in LD and SD environments with discontinuity in *DL*^s^ across the critical photoperiod range. This hinders investigation of the PRN-FTP for maize and accurate modeling of flowering time *G* × *E*, warranting further experimentation. The *DL*^s^ index can be used for the design of new MET networks to address this.

The PRN-FTP model provides an ecophysiological framework for decomposing and understanding phenotypic variation in flowering time measured in days. For a specific genotype, the pace of development towards reproductive transition is affected by temperature fluctuations within an environment, with the magnitude of this influence varying across environments based on the unique temperature patterns. When contrasting genotypes are present in the same environments, interactions between temperatures and genetic variation in rates of development (phyllochron) and durations of reproductive transition (BVP) gives rise to variation in the timing of photoperiod sensitization. In turn, genetic variation in critical thresholds and photoperiod sensitivities lead to secondary interactions with daily rates of photoperiodic change and exposure to further temperature fluctuations within and across environments. This highlights the interplay between genetic and environment effects that shape phenotypic variation in flowering time.

Lastly, as the PRN-FTP model is based on common ecophysiological processes, we expect that it can be extended to other crops. Investigating the trait space of PRN-FTP components could provide valuable insights into flowering time adaptations across species. Our findings underscore the significance of considering the historical context of adaptation and breeding in unraveling the genetic and physiological underpinnings of complex traits. This, in turn, holds promising implications for future crop improvement endeavors.

## Supporting information

Supporting lnformation

## Acknowledgements

This study was made possible thanks to the production and availability of datasets from prior and ongoing projects: the EXPOSE and PhotoGrid projects at INRAE-LEPSE (ANR-16-IDEX-0006; France 2030 program; Post AgreenSkills Fund), the INVITE project (European Union”s Horizon 2020 research and innovation program, grant no. 817970), the Maize ATLAS project (USDA-ARS, grant no. 2011-67003-30342), the Multiple Disease Resistance project (North Carolina State University, Cornell University, and University of Delaware collaboration), the Genomes-to-Fields Initiative (https://www.genomes2fields.org/home/), and the Panzea project (NSF-PGRP, grant no. 1238014). We wish to thank a number of people who contributed to this study: (i) Mainassara Abdou Zaman-Allah, Terence Molnar, and Prasanna Boddupalli from CIMMYT for providing much of the germplasm used for panel A; (ii) Alexis Bédiée, Llorenç Cabrera-Bosquet, Nicole Choquette, Arnaud Desbiez-Piat, Thomas Laisné, Noé Lalouette-Marrier d”Unienville, Maëlle Lis, Anthony Rosello, Javier Soto Mendoza, and Benoît Suard for supporting the PhenoArch platform experiments at Montpellier, FR; (iii) Arnaud Brunet, Bernard Lagardere, and Jean-Rene Loustalot from the experimental station at Saint-Martin-de-Hinx, FR for supporting germplasm management and field testing; (iv) the ISRA team Ibrahima Sarr, Malick Ndiaye, Awa Mbengue, Abdoulaye Hoth, and Alain Audebert (CIRAD) for field trial data from Nioro du Rip, SN; and (v) the Moreno Retis team José Moreno and Luis Rodríguez for field trial data from Puerto Vallarta, MX.

## Author Contributions

JD, RJW: conceptualization; CP, CW, RJW: resources; JD, CP, KJ, EM, RJW: investigation; JD, CP, JR, RJW: data curation; JD, PM, BP, RJW: methodology; JD, JR, KJ, RJW: formal analysis; JD, RJW: writing - original draft; EM, JR, PM, KJ: writing - review & editing; BP, CW, RJW: supervision; EM, RJW: funding acquisition.

## Data Availability

Data and metadata from this study are available in Supplementary Files at Dryad (https://doi.org/10.5061/dryad.x95x69pth). Sources of data are indicated in the methods sections and Table S1 (also see acknowledgments). Computer code for envirotyping *DL*^s^ and fitting the PRN-FTP is available at: https://github.com/maizeatlas/prnftp.

