## Supporting lnformation for "A reaction norm for flowering time plasticity reveals physiological footprints of maize adaptation"

The following Supporting Information is available for this article:

**Methods S1** Temperature response functions used to calculate thermal time.

**Methods S2** Description of additional experiments (platform and fields).

**Table S1** Summary of datasets used for this study.

**Figure S1** Comparison of the normalized root mean square error (NRMSE) for three different temperature response functions used to calculate thermal time.

**Figure S2** The distributions of sensed daylength ($DL^s$) for the 5th, 50th, and 95th percentile-maturity groups across environments in dataset D3.

**File S1** Estimates of thermal time from planting to male flowering (anthesis) for Genomes-To-Fields data used to compare temperature response functions.

**File S2** Estimates of thermal time to tassel initiation for panel 1 used for modeling sensed daylength ($DL^s$).

**File S3** Metadata for 680 field environments and estimates of sensed photoperiod (DLs) envirotypes.

**File S4** Data used for modeling the physiological reaction norm for flowering time plasticity (PRN-FTP) of seven genotypes.

**File S5** Data used for estimating parameters of the physiological reaction norm for flowering time plasticity (PRN-FTP) in a diversity panel constituting separate breeding pools of maize.

**File S6** Parameter estimates for the physiological reaction norm for flowering time plasticity (PRN-FTP) in a diversity panel constituting separate breeding pools of maize.

**Methods S1. Temperature response functions used to calculate thermal time.**

Three temperature response function used to calculate thermal time ($T_t$) were compared in their efficiency to model time to anthesis. The first equation assumes a segmental linear relationship between developmental rate and temperature integrated across time using daily minimum and maximum temperatures based on the SM(B,30-) formula according to Bonhomme *et al*. (1994):

$$f(T_X, T_N, T_b, T_o) = \begin{Bmatrix} \left[\frac{(T_X+T_N)}{2}\right] - T_b\,; & if\ T_X \le T_o\ \&\ T_N \ge T_b \\ \left[\frac{(T_X+T_b)}{2}\right] - T_b\,; & if\ T_X \le T_o\ \&\ T_N < T_b \\ \left[\frac{([T_o-(T_X-T_o)]+T_N)}{2}\right] - T_b\,; & if\ T_X > T_o\ \&\ T_N \ge T_b \\ \left[\frac{([T_o-(T_X-T_o)]+T_b)}{2}\right] - T_b\,; & if\ T_X > T_o\ \&\ T_N < T_b \end{Bmatrix} \qquad \text{(Eq. S1)}$$

where $T_X$ and $T_N$ are daily maximum and minimum temperatures (°C), respectively, $T_b$ is a base temperature fixed at 8 °C, and $T_o$ is an optimum temperature fixed at 30 °C. The $T_t$ was then calculated by accumulating the function by summation across days *i* to *n*.

The other two temperature response functions assume nonlinear relationships integrated across time using hourly temperatures. The first one was proposed by Wang *et al*. (1998), and was showed to accurately represent the rate of maize development (Wang *et al*. 2017). It is given by:

$$f(T_h, T_{min}, T_{opt}, T_{max,}\alpha, \beta) =$$

$$\begin{Bmatrix} 0\,; & if\ T_h \le T_{min} \\ \left[\frac{\left(2[T_h-T_{min}]^{\alpha}[T_{opt}-T_{min}]^{\alpha}\right)-(T_h-T_{min})^{2\alpha}}{\left(T_{opt}-T_{min}\right)^{2\alpha}}\right]^{\beta}\,; & if\ T_{min} < T_h < T_{max} \\ 0\,; & if\ T_h \ge T_{max} \end{Bmatrix},$$

$$\alpha = ln2/ln\left[(T_{max} - T_{min})/(T_{opt} - T_{min})\right],\ \beta = 1.5 \qquad \text{(Eq. S2)}$$

where, $T_h$ is hourly temperature (°C), $T_{min}$ is fixed at -17 °C, $T_{opt}$ is fixed at 31.5 °C, and $T_{max}$ is fixed at 43 °C. The value for $\beta$ was adapted for maize in order to minimize the difference in the shape of the response curve in relation to Parent & Tardieu (2012).

The third equation was proposed by Parent & Tardieu (2012), who used it to describe the rate of leaf development, and other developmental processes for several species.

$$f\left(A, T_h, \Delta H_A^{\ddagger}, R, \alpha, T_0\right) = A T_h e^{\left(\frac{-\Delta H_A^{\ddagger}}{R T_h}\right)} \Bigg/ 1 + \left[e^{\left(\frac{-\Delta H_A^{\ddagger}}{R T_h}\right)}\right]^{\alpha\left(1-\frac{T_h}{T_0}\right)}\,; \qquad \text{(Eq. S3)}$$

where, from Table S2 in Parent & Tardieu (2012), A is a scaling coefficient fixed at 5.16 x $10^{10}$, $T_h$ is hourly temperature (°K), $\Delta H_A^{\ddagger}$ (J $mol^{-1}$) and $T_0$ (°K) are shape parameters specific to each species (73900 J $mol^{-1}$ and 306.4 °K), R is the universal gas constant (8.314 J K-1), and $\alpha$ is a constant valid for most species fixed at 3.5 (dimensionless).

**Methods S2. Description of additional experiments (platform and field tests).**

***Phenotyping platform measurements of phyllohcron:*** An experiment was carried out on the Phenoarch platform (Cabrera-Bosquet *et al*. 2016) from January to April 2022 with 313 genotypes following a resolvable row-column design partitioned into six replicate blocks. The experiment was planted in two groups on January 19 and 20, 2022. Experimental units consisted of 9 L pots containing 30:70 (v/v) mixture of a clay and organic compost to which a single plant per genotype was assigned. Two check genotypes (B73 and CML341) were included with four replicates at fixed positions within each replicate block (i.e., a total of 24 plants per check genotype) to ensure a better coverage of the spatial variability. The design was generated using the DiGGer R package (Coombes 2009). Two levels of soil water content were imposed, either retention capacity (WW, mean soil water potential of -0.0015 MPa) or water deficit (WD) in which soil water potential was individually maintained at -0.23 Mpa for each pot after the majority of the plants reached six ligulated leaves. The greenhouse temperature was 25±3°C and 18±1°C during day and night. Air temperature and humidity were measured at six positions in the platform every 15 minutes (Prado *et al*. 2018). The temperature of the meristematic zone was also measured with fine thermocouples in 20 plants (one of the check genotypes). Supplemental light was provided either during daytime when external solar radiation was below 90 W $m^{-2}$ or to extend the photoperiod to 11 h with 0.4 lamps m-2.

The number of visible leaf tips of each plant was recorded every week using a decimal score, based on the proportional length of coverage by the youngest visible leaf compared to the subtending leaf (e.g., 7.0 indicated visibility of the $7^{th}$ leaf inside the whorl, while 7.5 indicated visibility of the $7^{th}$ leaf at 50% of the subtending leaf length). Phyllochron of each experimental unit (plant within a pot) was calculated by a simple linear regression of thermal time (Eq. S2) from seedling emergence on the number of visible leaves.

The current study used marginal means estimated only from the WW treatment to obtain genotype-specific values for phyllochron using a mixed linear model:

$$\boldsymbol{Y}_{ijklmnp} = \boldsymbol{\mu} + \boldsymbol{T}_i + \boldsymbol{K}_j + \boldsymbol{G}_{k(j)} + \boldsymbol{TK}_{ij} + \boldsymbol{TKG}_{ijk} + \boldsymbol{B}_{l(i)} + \boldsymbol{R}_{m(i)} + \boldsymbol{C}_{n(i)} + \boldsymbol{f}(\boldsymbol{r},\boldsymbol{c})_{p(i)} + \boldsymbol{\varepsilon}_{jklmnp(i)} \;;$$

where, $\boldsymbol{Y}_{ijklmnp}$ is the pot specific plant phyllochron; $\boldsymbol{\mu}$, $\boldsymbol{T}_i$, $\boldsymbol{K}_j$, and $\boldsymbol{G}_{k(j)}$ are fixed effects of the overall mean, the $i^{th}$ treatment, $j^{th}$ check group, and $k^{th}$ genotype nested in the $j^{th}$ check group, respectively; $\boldsymbol{TK}_{ij}$, $\boldsymbol{TKG}_{ijk}$, $\boldsymbol{B}_{l(i)}$, $\boldsymbol{R}_{m(i)}$, $\boldsymbol{C}_{n(i)}$ are random effects of the interaction between the $i^{th}$ treatment and $j^{th}$ check group, interaction between the $i^{th}$ treatment and $k^{th}$ genotype nested in the $j^{th}$ check group, the $l^{th}$ replication nested in the $i^{th}$ treatment, the $m^{th}$ row nested in the $i^{th}$ treatment, and the $n^{th}$ column nested in the $i^{th}$ treatment, respectively; $\boldsymbol{f}(\boldsymbol{r},\boldsymbol{c})_{p(i)}$ is a random effect of the $p^{th}$ row-column spatial kernel nested in the $i^{th}$ treatment; $\boldsymbol{\varepsilon}_{jklmnp(i)}$ is the residual nested in the $i^{th}$ treatment. The mixed linear model was fit using the R package *Sommer* (Covarrubias-Pazaran 2016).

***Phenotyping platform measurements of tassel initiation:*** An experiment was carried out on the PhenoArch platform (Cabrera-Bosquet *et al*. 2016) from January to April 2024 in order to estimate tassel initiation from temporal dissections of the shoot apical meristem (SAM). The experiment was planted on January 26, 2024. All plants were grown in the same platform environment with day / night temperature conditions of 23.9±2°C / 18.4±1°C, respectively, and a natural photoperiod supplemented with artificial light when external solar radiation was below 90 W $m^{-2}$ or to extend the photoperiod for maintaining an 11 h daylength (from 7:30 to 18:30).

A factorial design with a light treatment and genotype treatment was used. There were two levels of the light treatment, which included a 30s nightbreak, and a no-nightbreak control. Beginning two-weeks after sowing, and continuing for three weeks, the nightbreak treatment was applied for 30s within imaging units of the platform equipped with LED illumination (5050–6500 K colour temperature) between 21h00 and 23h00. Ten levels of the genotype treatment were used, which included inbred lines 2369, B73, CML258, CML277, CML341, CML373, LH123Ht, Tzi8, and Tzi9 (in addition to an undetermined inbred line due to a seed stock error).

Experimental units consisted of 9 L pots containing 30:70 (v/v) mixture of a clay and organic compost to which a single plant per genotype was assigned to a structured layout. The pot layout was arranged as 10 rows x 48 columns in the platform. Light treatments (30s nightbreak vs. no-nightbreak control) were assigned to pots in alternating rows. Twelve sampling blocks, used for SAM dissections, were defined in sets of four columns containing two replicate plants of each genotype. The DiGGer R package (Coombes 2009) was used to randomize genotypes across sampling blocks per light treatment.

For dissection and imaging of the SAM, destructive sampling of the sampling blocks occurred sequentially, beginning four weeks after sowing and continuing every three days over the course of three weeks until all genotypes reached tassel initiation. The number of visible and ligulated leaves, maximum length and width of each leaf rank, collar height of each leaf rank, and distance between collars was also measured for each plant. The SAM was dissected and imaged with a Leica EZ4 dissecting microscope (Wetzler, Germany). Development of the SAM was scored by visual inspection using a customized scoring scale (1-10) informed by Bonnett (1966) and Krüger (1984), spanning pre-elongation of the SAM to final stages of tassel formation. Tassel initiation corresponded to the point at which tassel-branch primordia were apparent. Scores were significantly ($p < 0.05$) and highly correlated ($r = 0.9$) with the length:width ratio of the SAM, measured using ImageJ. Tassel initiation inferred from visual scoring was consistent with a SAM length:width ratio of 2.0, determined by linearly regressing the length:width ratio to the number of visible leaf tips at the time of dissection. The current study used only the data from the no-nightbreak control treatment.

***Field measurements of final leaf number*:** In 2023, a field experiment was carried out at Puerto Vallarta, MX, planted on January 9, 2023. Experimental units consisted of 480 plots (3 m length) with an interplant spacing of 25 cm. Plots were laid out in 40 rows x 12 ranges with an interplot spacing of 75 cm, partitioned into two sections of 4 ranges (160 plots) and 8 ranges (320 plots). The trial was surrounded by border plots to minimize edge effects. The DiGGer R package (Coombes 2009) was used to apply a resolvable row-column p-rep design to each section, which contained 114 and 256 genotypes, respectively. Section 1 included 92 unreplicated genotypes, two replicates of 18 genotypes, and four replicates of eight genotypes. Section 2 included 192 unreplicated genotypes, two replicates of 64 genotypes replicated twice. Three of the same genotypes were present in both sections, such that a total of 367 unique genotypes were tested.

Data were collected based on assessment of each plot (emergence date) or six individual plants per plot (leaf number and flowering time). Emergence dates were visually assessed and recorded as the first sign of emergence from the soil surface. Leaf numbers of individual plants were tracked by tagging the 5th, 10th, and 15th leaf during development, followed by recording of final leaf number. Female (silking) and male (anthesis) flowering dates of individual plants were recorded as the first sign of silks and anthers, respectively.

The data were analyzed using the mixed model software Echidna (Gilmour 2021). Days to emergence, final leaf number, and days to flowering were modeled as:

$$\boldsymbol{Y}_{ijkl} = \boldsymbol{\mu} + \boldsymbol{\gamma}_i + \boldsymbol{B}_j + \boldsymbol{R}_k + \boldsymbol{C}_l + \boldsymbol{\varepsilon}_{ijkl} ;$$

where, $\boldsymbol{Y}_{ijkl}$ are the phenotypic values for the given trait; $\boldsymbol{\mu}$ and $\boldsymbol{\gamma}_i$ are fixed effects of the overall mean and $i^{th}$ genotype, respectively; $\boldsymbol{B}_j$, $\boldsymbol{R}_k$, and $\boldsymbol{C}_l$ are random effects of the $j^{th}$ block, $k^{th}$ row, and $l^{th}$ range, respectively; $\boldsymbol{\varepsilon}_{ijkl}$ is the residual. Random effects were assumed to be normally distributed $\mathbf{B} \sim N(0, \sigma_b^2)$, $\mathbf{R} \sim N(0, \sigma_r^2)$, $\mathbf{C} \sim N(0, \sigma_c^2)$, and $\boldsymbol{\varepsilon} \sim N(0, \sigma_\varepsilon^2)$. For each of the fitted models, a likelihood ratio test was used to select the most parsimonious model for the specific trait and environment. Spatial modeling of the row and column residual variation was tested but did not improve the model fit based on the Bayesian information criterion.

**Table S1. Summary of datasets used in this study.**

| Dataset | Objective | Projects[a] | Experiment | Number of genotypes[b] | Number of environments[c] | Number of datapoints | Supplemental files |
|---|---|---|---|---|---|---|---|
| D1 | Select thermal time model | G2F | Field | 24 | 25-26 | 614 | S1 |
| D2 | Estimate genotype-specific BVP (ºCd) | EXPOSE | Platform | 284 | 1 | 284 | S2 |
| D3 | Envirotype $DL^s$ | G2F; Invite; Maize ATLAS; PhotoGrid; MDR; PhotoGrid | Field | 3 | 671 | 2013 | S3 |
| D4 | Model PRN-FTP | Maize ATLAS; PhotoGrid; MDR; Panzea | Field | 14 (7 tropical; 7 temperate) | 13-37 | 351 | S4 |
| D5 | Trait space analysis | MDR; Panzea | Field | 281 | 4-19 | 4141 | S5; S6 |

[a]Data from different projects were used for this study. New data contributed specifically by this study were from projects EXPOSE (platform experiments) and PhotoGrid (field experiment).
[b]Dataset D5 included 281 genotypes. The results are provided for all of the genotypes in the supplemental files. In the main text, a subset of 236 genotypes classified as temperate, tropical, and admixed were used.
[c]The range indicates the minimum and maximum number of environments among the genotype.

**Figure S1** Comparison of the normalized root mean square error (NRMSE) for three different temperature response functions used to calculate thermal time.

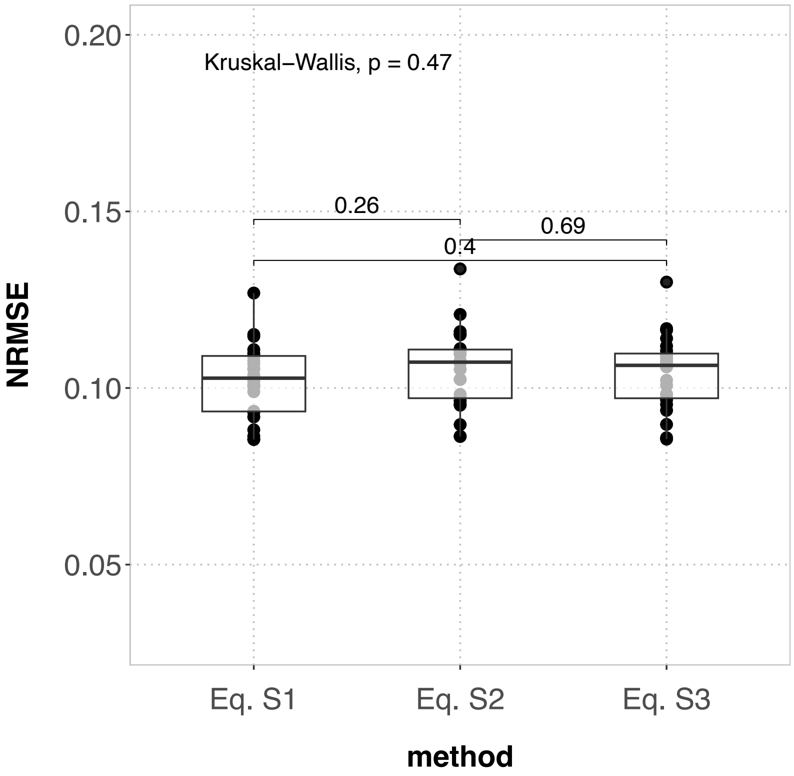


Boxplots of the NRMSEs for thermal time from planting to male flowering (anthesis) estimated for 24 hybrids based on equations S1-S3 (Methods S1).

**Figure S2** The distributions of sensed daylength ($DL^s$) for the 5$^{th}$, 50$^{th}$, and 95$^{th}$ percentile-maturity groups across environments in dataset D3.

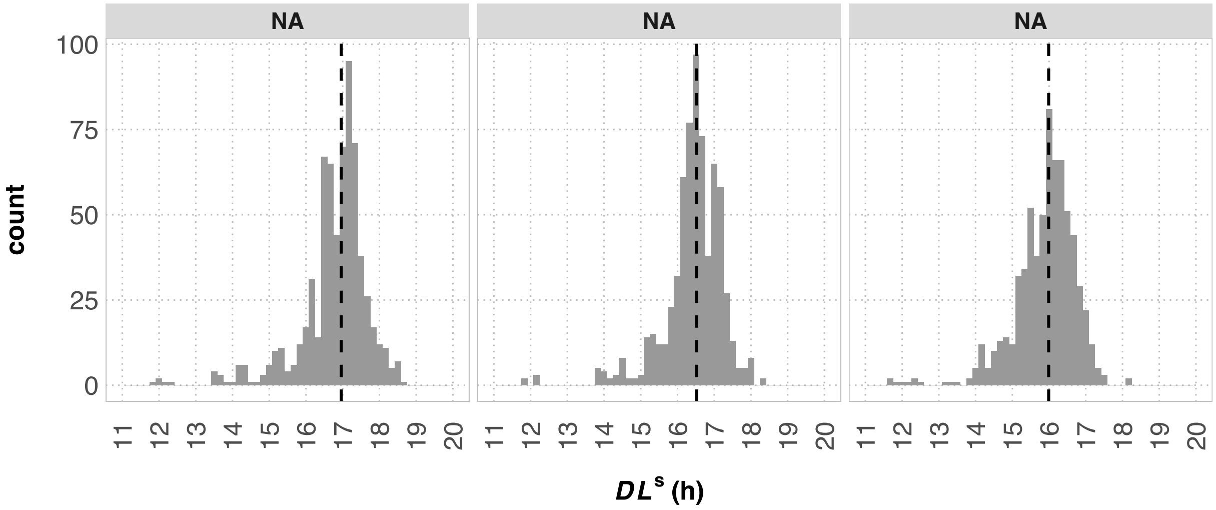


Histograms of the $DL^s$ distribution for the 5$^{th}$, 50$^{th}$, and 95$^{th}$ BVP percentile-maturity groups across 671 field environments (File S3). The median $DL^s$ of each percentile-maturity group is indicated by the dashed line (17.0 h, 16.5 h, and 16.0 h for the 5$^{th}$, 50$^{th}$, and 95$^{th}$ percentile groups respectively).
